# Common and distinct roles of frontal midline theta and occipital alpha oscillations in coding temporal intervals and spatial distances

**DOI:** 10.1101/2020.08.05.237677

**Authors:** Mingli Liang, Jingyi Zheng, Eve Isham, Arne Ekstrom

## Abstract

Judging how far something is and how long it takes to get there are critical to memory and navigation. Yet, the neural codes for spatial and temporal information remain unclear, particularly the involvement of neural oscillations in maintaining such codes. To address these issues, we designed an immersive virtual reality environment containing teleporters that displace participants to a different location after entry. Upon exiting the teleporters, participants made judgements from two given options regarding either the distance they had travelled (spatial distance condition) or the duration they had spent inside the teleporters (temporal duration condition). We wirelessly recorded scalp EEG while participants navigated in the virtual environment by physically walking on an omnidirectional treadmill and traveling through teleporters. An exploratory analysis revealed significantly higher alpha and beta power for short distance versus long distance traversals, while the contrast also revealed significantly higher frontal midline delta-theta-alpha power, and global beta power increases for short versus long temporal duration teleportation. Analyses of occipital alpha instantaneous frequencies revealed their sensitivity for both spatial distances and temporal durations, suggesting a novel and common mechanism for both spatial and temporal coding. We further examined the resolution of distance and temporal coding by classifying discretized distance bins and 250ms time bins based on multivariate patterns of 2-30 Hz power spectra, finding evidence that oscillations code fine-scale time and distance information. Together, these findings support partially independent coding schemes for spatial and temporal information, suggesting that low-frequency oscillations play important roles in coding both space and time.

## Introduction

### Background

Tracking where we are in space and time is important for both navigation and episodic memory (Eichenbaum & Cohen, 2014; Ekstrom & Isham, 2017; Robin & Moscovitch, 2014; Tulving, 2002). However, it is not clear what neural mechanisms are recruited for spatial and temporal coding in humans and whether they share similar coding principles (Ekstrom & Isham, 2017; Frassinetti et al., 2009; Walsh, 2003). Movement, either physical or imagined, is a core part of both our sense of space and time, and induces robust hippocampal low-frequency oscillations (3-12Hz) in both rats (Vanderwolf, 1969) and humans (Bohbot et al., 2017; Ekstrom et al., 2005; Goyal et al., 2020; Jacobs, 2013; Watrous et al., 2011). Because movement typically involves changes in both space and time, one possibility is that low-frequency oscillations play a role in coding both variables.

Past investigations have established an important role for hippocampal theta oscillations in coding spatial distance in humans but evidence is lacking for the role of neocortical theta oscillations in distance coding. For example, hippocampal theta power increases linearly with the amount of distance travelled in virtual reality (Bush et al., 2017; Vass et al., 2016), cross-regional theta connectivity plays a critical role in judgments of relative spatial distance (Kim et al., 2018), and theta network connectivity differentiates distance from temporal contextual retrieval (Watrous et al., 2013). However, it is not clear whether neocortical theta oscillations can code spatial distance in a similar fashion, and if scalp EEG can reveal such a cortical theta distance code.

In addition, while past studies have established a role for low-frequency oscillations in spatial distance coding, their role in representing temporal durations remains less clear. The medial temporal lobes (MTL) of rodents are capable of internally generating representations that track time passage (Itskov et al., 2011; MacDonald et al., 2011; Pastalkova et al., 2008; Wang et al., 2015). Given the strong presence of delta and theta oscillations in MTL, it is possible that low-frequency oscillations contribute to temporal duration coding and that such a time code can manifest in neocortical low-frequency oscillations as well. Past studies have also revealed a role for cortical beta oscillations in supporting duration reproduction in humans, such as the finding that increased alpha-beta coupling strengths yield better time reproduction precision (Grabot et al., 2019), and higher beta power recorded with scalp EEG predicts longer reproduced durations (Kononowicz & van Rijn, 2015). Therefore, both delta-theta and beta band oscillations are strong potential candidates specifically dedicated to temporal duration coding, or both spatiotemporal coding, an issue we seek to resolve here. Beside low-frequency power changes, another possible oscillatory timing mechanism is alpha frequency modulation. Alpha frequency variations manifest independently of changes in alpha power (Samuel et al., 2018), and alpha frequency modulation has been implicated in the temporal resolution of visual perception (Cecere et al., 2015; Samaha & Postle, 2015). Nonetheless, how alpha frequency fluctuations relate to duration timing remains unclear and unresolved.

### Objectives

In this current study, we aim at experimentally dissociating the spatial distance and temporal duration information available to participants. Then, we examine whether and how low-frequency oscillations support spatial distance and temporal duration coding, and whether such spatiotemporal processing shares similar coding schemes. To address these research questions, we developed a teleportation task in an immersive and ecologically enriched virtual environment (Figure 1), largely similar to the experimental design in Vass et al. (2016) and capable of disentangling spatial and temporal information. In this task, participants entered a virtual teleporter, were presented with a black screen for a couple of seconds, and then exited at a different location in the virtual environment. After exiting, participants were prompted to make a binary-choice judgment regarding the distance they were transported inside the teleporter (the spatial distance task) or how long the duration was they spent inside the teleporter. By manipulating the distance and duration information independently, we disentangled participants’ memory for spatial distance from that of temporal duration. This in turn allowed us to examine their neural correlates separately. In addition, participants navigated around the virtual reality by physically walking on an omnidirectional treadmill while wearing a head mounted display, allowing us to study the relationship between cortical oscillations and spatiotemporal processing under more ecologically enriched conditions.

**Figure 1.**
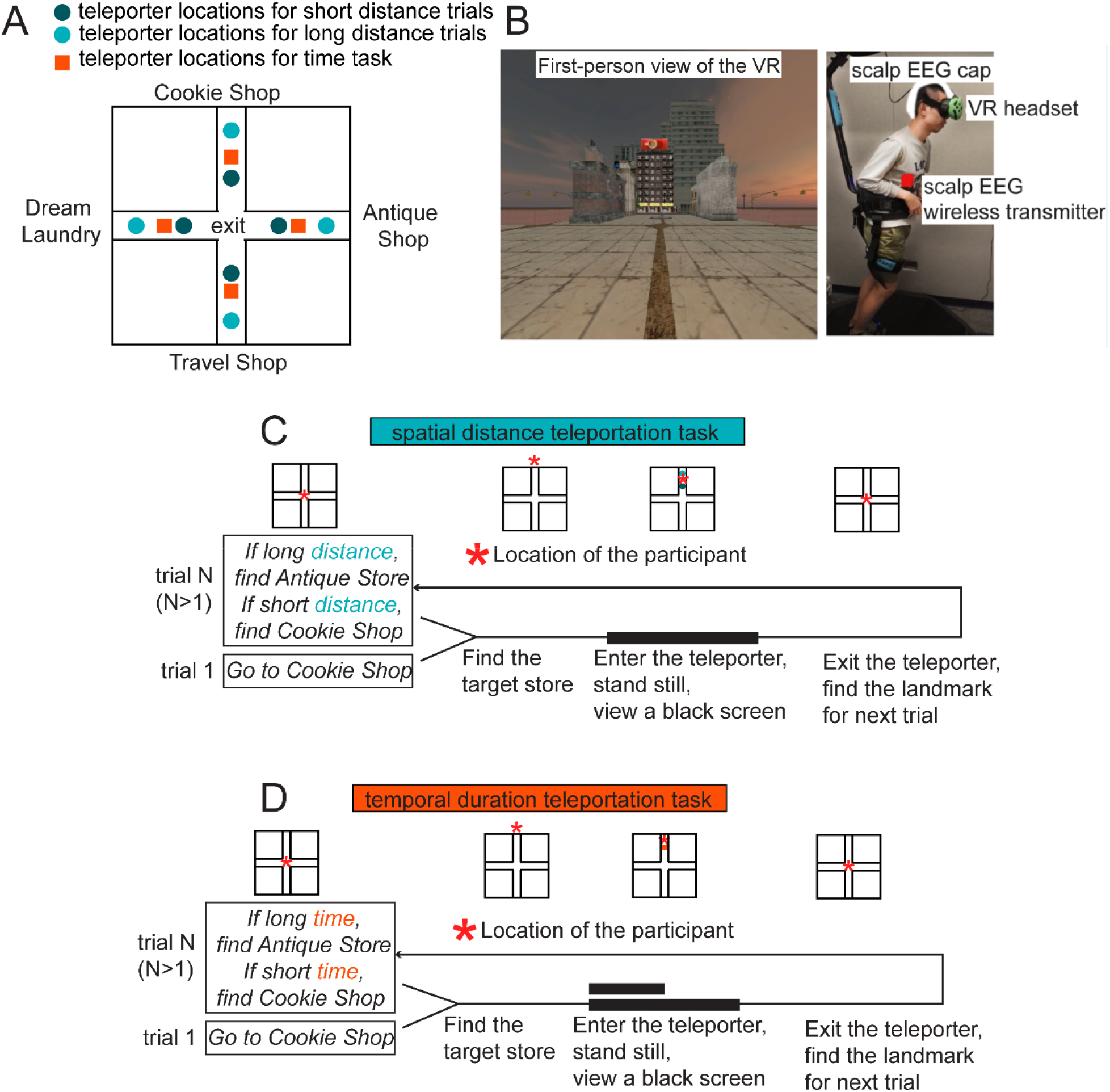
Spatial and temporal teleportation tasks, and virtual reality (VR) setup. **(A)** Layout of the VR and the possible entry locations of teleporters. **(B)** A view of the virtual environment, and the VR-scalp EEG setup. **(C)** Task flow in the spatial task. Participants were either teleported a short or long distance inside teleporters. **(D)** Task flow in the temporal task. Participants either experienced a short (4 seconds) or long (8 seconds) duration inside teleporters while standing still.

### Hypotheses

We tested two primary hypotheses. First, for the **within-task difference hypothesis**, we tested whether cortical oscillatory power (2-30Hz) and occipital alpha frequencies responded differently within tasks, i.e., judging short vs. long spatial distance, or short vs. long temporal durations. Second, for the **between-task difference hypothesis,** we tested whether such oscillatory codes differed between tasks, i.e. for spatial distance vs. temporal duration judgments, which might further support the ideas of independent codes (Watrous et al., 2013) vs. a common magnitude estimation mechanism (Walsh, 2003) for spatiotemporal coding. Together, these analyses allowed us to address to what extent the coding for spatial distance and temporal durations involves common vs. distinct neural mechanisms.

## Materials and Methods

This study was approved by the Institutional Review Board at the University of Arizona, and all participants provided informed consent. The data analyzed in this study are available at https://osf.io/3vxkn/.

### Participants

We tested 19 adults (7 females, 12 males) from the Tucson community. Because this is the first investigation of its type (scalp-recorded oscillatory correlates of spatiotemporal processing), it is difficult to estimate exact effects sizes needed to determine the sample size. Therefore, we based our sample size on a previous study in which we observed movement-related changes in low-frequency oscillations during navigation (Liang et al., 2018). Participants received monetary ($20/hour) and/or class credit for compensation. Prior to testing, participants received a virtual reality training session, which involved 30 minutes of walking on the omnidirectional treadmill with a head mounted display on. We implemented the training to screen out participants with potential susceptibility to cybersickness.

### Stimuli, Apparatus and Virtual Reality

The virtual environment was constructed with the Unity Engine and rendered with an HTC Vive headset. Immersive walking experiences were simulated with an omnidirectional treadmill (KAT VR Gaming Pro, KAT VR, Hangzhou China). Physical walking motions on the omnidirectional treadmill were translated into movements in the virtual reality.

The size of the virtual environment was 560 × 560 virtual square meters. The layout of the virtual environment was a plus (+) sign (Figure 1A), with four arms extending from the center. Four target stores were placed at the end of each arm (Cookie Shop, Dream Laundry, Antique Store, and Travel Shop). Identical filler buildings were placed along each arm.

The entry point to the teleporters was rendered as a purple circle. When participants “collided” with teleporters in the virtual reality, they initiated a teleportation event. During teleportation, they stood still for a few seconds while viewing a black screen on the head-mounted display, and eventually exited at the center of the plus maze.

### Behavioral Tasks

Participants completed two tasks: a spatial distance task and a temporal duration task. In the spatial task, the teleporters displaced the participants with one of the two possible spatial distances while the teleportation duration was kept constant. In the temporal task, the teleportation process could last a short (4 seconds) or long (8 seconds) duration, while the teleporters transported the participants a fixed distance. Each task involved 48 trials.

### Navigation phase

At the beginning of a trial, participants started at the center of the plus maze and navigated to a target store. The target store was either specified for the first trial, or it needed to be determined for the following trials. When arriving at the target store, participants entered a dummy teleporter in front of the target store. This involved showing a black screen for 4 seconds and rotating participant’s camera angle by 180 degrees. This dummy teleporter was set up to timestamp participants’ arrival times on the EEG and was not used in any subsequent analyses. If participants arrived at the wrong store, the dummy teleporters sent participants back to the center of the plus maze and they searched for the store again. During the navigation phase, no teleporters were visible except for four dummy teleporters in front of four target stores to detect arrivals at the correct store.

### Teleportation phase

After navigating to the target store, participants then walked up to and entered a new teleporter spawned in front of the target store. In the spatial distance task, for long distance trials, the teleporters spawned 200 virtual meters away from the center of the plus maze, and for short distance trials, the teleporters spawned 100 virtual meters away from the center. In the temporal duration task, the teleporters spawned 144 meters away from the center. Upon entering the teleporter, participants stood still, with the camera fading to a completely black screen in 200 milliseconds. They viewed the black screen for a specific duration (spatial task: 5.656 seconds, and temporal task: 4 or 8 seconds). Then participants reemerged at the center of the plus maze, with their camera fading from pure black to the view standing at the center of plus maze, in 200 milliseconds.

### Judgment phase

After exiting the teleporter, written instructions were provided to the participants by showing a billboard message overlaid on top of the virtual reality view. The instructions were used to decide which target store to visit for the current trial. For the spatial task, instructions were: “*If far distance, go find store A. If short distance, go find store B.*” For the temporal task, instructions were: “*If long time, go find store A. If short time, go find store B.*” The instructions in the virtual reality disappeared when participants walked further than 55 meters away from the center of the plus maze. By asking participants to judge spatial distance and temporal durations, we ensured that they maintained these two task-relevant variables.

### Parameters for the behavioral tasks

For the spatial task, the duration of viewing the black screen was 5.656 seconds for both long distance and short distance trials. Short distance was defined as teleporting 100 meters and long distance was defined as teleporting 200 meters and (Figure 1C). For the temporal task, the distance teleported was kept constant, at 141.4 meters. For short duration trials, participants viewed 4 seconds of a black screen during teleportation, while for long duration trials, they viewed 8 seconds of a black screen (Figure 1D). We selected these parameters for our spatial and temporal tasks to ensure the average teleportation speeds were the same between spatial and temporal tasks: the average teleportation speed for the spatial task was 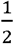 x (200 meters/5.656s + 100 meters /5.656s) ≈ 26.52 m/s while the average speed for the temporal task was 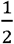 x (141.4 meters/8s + 141.4 meters/4s) ≈ 26.51 m/s). This is because movement speed has been shown to affect low-frequency oscillations (Caplan et al., 2003) and thus we attempted to control for movement speed during teleportation.

The order of short/long trials were pseudorandomized across the 48 trials. Short and long teleportation each had 24 trials, with each target store visited 12 times. Two sets of short/long orders were generated so that spatial and temporal tasks did not use the same set of long/short sequences. The order of task types, and the short/long sequence sets, were counterbalanced across participants. Before starting the main experiment, participants were shown three examples each: long distance teleportation, short distance teleportation, short temporal duration teleportation, and long temporal duration teleportation. Some participants repeated this practice procedure until they reported understanding the differences between short/long trials.

After each block of 12 trials, participants had the option to take a short break of 3 minutes. When participants took a break, we first asked participants to stand still and relax for 90 seconds on the omnidirectional treadmill while wearing the head-mounted display and viewing a black screen. Then we recorded the 90 second EEG data as the baseline. Pooling across the spatial and temporal tasks, we recorded, on average, 364.74 seconds (SD: 183.64 seconds) of EEG baseline data.

### EEG Acquisition and Preprocessing

The continuous EEG was recorded with a 64-channel BrainVision ActiCAP system, which included a wireless transmission MOVE module, and two BrainAmp amplifiers (BrainVision LLC, Morrisville, NC). We recorded from 64 active electrodes, placed on the scalp according to the international 10-20 system. The reference electrode was located at FCz, and no online filter was applied to the recordings. Before the experimenter proceeded to start the recordings, impedances of all 64 electrodes were confirmed below 5kΩ.

Preprocessing and analyses were performed with EEGLAB (Makeig et al., 2004), and customized codes in MATLAB. No offline re-referencing or interpolation of electrodes was performed on the continuous data. A 1650^th^ order Hamming windowed sinc finite impulse response (FIR) filter was performed for 1-50Hz bandpass filtering on the continuous data using the EEGLAB pop_newfilt() function, with a transition bandwidth of 1 Hz, the passband edges of 1 and 50 Hz, and cut-off frequencies (−6dB) of 0.5 and 50.5 Hz. Artifact subspace reconstruction (ASR) was then applied to the filtered continuous data, with the EEGLAB *clean_asr()* function, to repair large amplitude spikes that were 5 standard deviations away from the clean segments of the continuous data.

### EEG Epoching and Segmentation

The continuous EEG data were segmented using a time window aligned with the start and ends of teleportation (not including the fade-to-black or fade-to-clear 200ms windows). This segmentation procedure yielded 48 epochs with a length of 5.656 seconds for the spatial task, and 48 epochs with a length of either 4 or 8 seconds for the temporal task. No baseline correction was applied. To keep the number of trials constant across participants, we did not reject trials based on incorrect behavioral responses. We did not reject trials based a voltage threshold because we mainly used independent component analysis to correct artifacts, as described below.

### Independent Component Analysis

Independent component analysis (ICA) with the infomax algorithm was performed in EEGLAB to correct artifacts. Note that we ran ICA on the artificial “continuous data structure” by concatenating all the data in the distance task, time task, and the resting baseline task. Our motivation was data in those three tasks should receive identical ICA correction procedure. We used an automatic component selection procedure, ICLabel (Pion-Tonachini et al., 2019) to avoid experimenter bias in identifying noisy components. Components were rejected automatically if they had labels of “Muscle”, “Eye”, “Heart”, “Line Noise”, or “Channel Noise” if their probability was higher than 90% for being one of those labels. On average, 8.84 (13.81% of all components, SD: 3.91) components were rejected.

### Time Frequency Analysis

#### Power measures for delta, theta, alpha, and beta bands

We estimated the instantaneous power during the teleportation windows with 6-cycle Morlet Wavelets using code from Hughes, Whitten, Caplan, and Dickson (2012). We sampled frequencies from 2 to 30 Hz in 20 logarithmic frequency steps, i.e., 2 Hz, 2.31 Hz, 2.66 Hz, 3.07 Hz, and 3.54 Hz for delta band, 4.08 Hz, 4.70 Hz, 5.42 Hz, 6.25 Hz and 7.21 Hz for theta band, 8.32 Hz, 9.59 Hz, and 11.06 Hz for alpha band, 12.76 Hz, 14.71 Hz, 16.96 Hz, 19.56 Hz, 22.56 Hz, 26.01 Hz, and 30 Hz for beta band. Zero paddings were added to both ends of the signal to alleviate edge artifacts. No baseline correction was applied to the power estimates. Logarithmic transform with a base of 10 was applied to the obtain power values before averaging. Mean power for each band was measured as log power averaged across timepoints within the teleportation window, across frequencies within a band, and across trials of interest.

#### Cluster-based permutation tests for multiple comparison correction

Cluster-based permutation tests (Maris & Oostenveld, 2007) were used to determine the statistical significance between the mean power values for short vs. long trials. Correction for multiple comparisons was implemented in Fieldtrip. First, to identify uncorrected significant samples, 64(electrodes)*4(frequency bands) = 256 Wilcoxon signed rank two-tailed tests were performed for the power contrasts, alpha = 0.05. Clusters were found by connecting significant sample pairs (electrode x frequency bands) with spatiospectral adjacency (minimum neighbor of channels was set to 0), and cluster-level statistics were computed using a weighted-sum (Hayasaka & Nichols, 2004) of all the z values returned by Wilcoxon signed rank tests within a cluster. Second, a surrogate distribution of cluster-level statistics was generated by randomly shuffling condition labels 1000 times on the subject level and retrieving the maximum cluster-level test statistic for each permutation. Third, p values of the observed cluster statistics were obtained by benchmarking to the surrogate distribution. Empirical clusters with a p value smaller than 0.025 (either left tail or right tail) were be reported.

We chose the nonparametric Wilcoxon signed rank tests over the parametric paired t tests because the normality assumption for t tests was violated. For all the power spectra contrast we conducted, all the power spectra differences showed a distribution different from normal distributions (one sample Kolmogorov-Smirnov test, alpha = 0.05, all p’s < 0.01). In the results reported in which we employed the Wilcoxon signed rank tests, medians instead of means were reported.

#### Effect sizes calculation

Cohen’s d was used as an estimate for effect sizes. For a within-participant paired comparisons between condition 1 and condition 2, we estimated the effect sizes using the following formula:

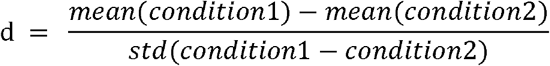

#### Frequency measures for alpha (8-12 Hz) band

To estimate alpha frequency, we used a frequency sliding technique (Cohen, 2014) to estimate the alpha frequency fluctuations. We first used a 125^th^ order finite impulse response (FIR) 8-12 Hz bandpass filter (using MATLAB firls() function) on the segmented EEG data, with a transition bandwidth of 1.2 and 1.8 Hz, the passband edges of 8 and 12 Hz, and cut-off frequencies (−6dB) of 7.12 and 12.98 Hz. We then employed the Hilbert Transform on the filtered segmented EEG data to obtain the instantaneous phase estimates of alpha oscillations during teleportation windows. Instantaneous frequencies at timepoint *t* were estimated as

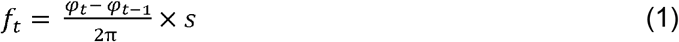

where *f* is the estimated instantaneous alpha frequency, *φ* is the estimated phase, and *s* is the EEG sampling rate. Here, we defined and estimated the instantaneous frequencies based on how many cycles the phase of alpha oscillations could go through in 1 second. Then, to smooth the frequency estimates, we applied a 10^*th*^ order median filter. We dropped the frequency estimates for the first 100ms and last 100ms for every trial because of potential inaccurate estimates of frequencies at the edges of signal.

We selected the following occipital electrodes to analyze their alpha frequency based on two criteria: visible alpha prevalence in the raw traces and an identical cluster of occipital electrodes to what we used in our past study (Liang et al, 2018). These 18 electrodes corresponded to: Pz, P3, P7, O1, Oz, O2, P4, P8, P1, P5, PO7, PO3, POz, PO4, PO8, P6, P2, and Iz.

Alpha frequency for each behavioral task is measured as alpha frequency estimates averaged across timepoints during the windows of interest, averaged across electrodes of interest, and averaged across trials of interest. To compare the alpha frequency variations between two conditions, we submitted the averaged alpha frequencies of 19 participants to two-tailed Wilcoxon signed rank tests (alpha = 0.05.) Six Wilcoxon signed rank tests were conducted, and the p values reported in the results section were FDR corrected (Benjamini & Yekutieli, 2001; Groppe, 2021), with the false discovery rate set to 0.05.

### Classification Analyses

#### Binary classification of the duration/distance types

To further confirm the role of frontal midline delta-theta oscillations in spatial and temporal judgments, a binary support vector machine (SVM) classifier was used to decode the types of teleportation using power of delta, theta, alpha and beta band, averaged at specific electrodes. For delta power, theta power, and alpha power, 4 electrodes around frontal midline region were selected (Fz, FC1, Cz, FC2.) For beta power, all available electrodes (64 electrodes) were chosen. Binary SVM classifiers were implemented in MATLAB, with the function *fitcsvm()*, with the kernel function set up as linear. Three decoding tasks on a within-subject level were implemented: 1) decoding whether the trial was from the teleportation trials that travelled short or long distance, 2) decoding whether the trial was from short duration trials, or the 4-8second portions of long durations trials in the time task, and 3) decoding whether the trial was from short duration trials or the 0-4 second portions of long durations trials. The ratio of train-test split for each iteration was 67%-33%. The training-testing sampling procedure was reiterated 1000 times for each participant, and for each decoding task. An accuracy percentage score was calculated using the predicted and actual labels of the testing data. The final decoding accuracy scores for 19 participants were submitted to two-tailed Wilcoxon signed rank tests, against the null hypothesis that the decoding accuracy was 50%. In total, 12 tests were conducted in the binary classification analysis, and the p values were FDR corrected (Benjamini & Yekutieli, 2001; Groppe, 2021), with the false discovery rate set to 0.05.

Additionally, we implemented a between-task classifier (space vs. time tasks) on an intersubject level. We combined trials from the space task and the time task across 19 participants, resulting in a dataset of 912 trials. Then, we tested whether we could successfully decode the task labels using the 912-trial dataset. By performing the classification on an intersubject level (with the task orders were counterbalanced), we avoided the possible confound of systematic drift over the course of experiment, which could have affected our decoding accuracy due to the blocked nature of the spatial vs. temporal judgments in our design (Benwell et al., 2019). For features used for training classifiers, we employed the 2-30 Hz power spectra from 64 electrodes averaged within each trial, resulting in 20*64 = 1280 features. The ratio of train-test split for each iteration was 67%-33%. The train-test split was repeated 100 times. To determine the statistical significance of decoding accuracy, we submitted the accuracies from 100 iterations to a two-tailed Wilcoxon signed-rank test against the null hypothesis of 50%.

#### Fine-scale time decoding analyses

To examine whether continuous time codes were present in the scalp EEG signal, SVM classifier was trained to decode times beginning at the onset of teleportation using the 2-30 Hz power spectra from 64 electrodes. The SVM algorithm was implemented in MATLAB using the *fitcecoc*() function, with coding style as ‘*onevsall’*, and other parameters as default.

250-millisecond timebins were extracted by discretizing 2-30 Hz power estimates. The size of timebins was chosen as the same one used by Bright et al. (Bright et al., 2020). Therefore, short/long distance teleportation trials (5.656 seconds) yielded 22 bins (22 * 250ms = 5.5 seconds, the last 156ms of data were dropped), short temporal duration trials (4 seconds) yielded 16 bins, and long temporal duration trials (8 seconds) yielded 32 bins. For the resting baseline data (90 seconds long for each resting session), we broke 90 seconds into continuous segments of 4 seconds, and from there, each 4s of baseline data were segmented into 16 bins.

Power estimates within each time bin were averaged over time, and the resulting power spectra within each bin were used to trained classifiers. The number of features were 20 frequencies x 64 electrodes = 1280 features. For each classification iteration, train-test split ratio was 75%-25%. To increase the independence between training sets and testing sets, a consecutive block of trials was reserved as the testing data, and the rest of data was used for training. Given our way of splitting the data, we were able to reiterate the classification procedure limited number of times: for the distance task, the procedure was repeated 37 times; for the short interval and long interval trials, 19 times; and for the baseline task, 16 times.

We calculated the accuracy score by summing how many correct predictions were made in 100 iterations for each timebin label. The accuracy scores were then averaged across all iterations yielding a final accuracy score for each participant. Given that number of timebins were different across the distance task, time task and baseline task, comparisons between them would be difficult. We standardized the accuracy scores as the accuracy ratios by dividing them against the chance level performance 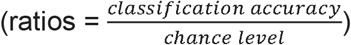. For the distance task time decoder, the chance level was 1/22 = ~4.55%; for decoding time in short temporal duration trials, the chancel level was 1/16 = 6.25%; for decoding time in long duration trials, the chance level was 1/32 = 3.125%, and for decoding time in the baseline data, the chance level was 1/16 = 6.25%.

To test whether we successfully decoded fine-scaled temporal information above chance, we submitted the standardized accuracy ratios for 19 participants to a two-tailed Wilcoxon signed rank tests against the null hypothesis that the accuracy ratios were different from 1. Ten signed rank tests were performed for this hypothesis, and the p values were FDR corrected (Benjamini & Yekutieli, 2001; Groppe, 2021), with the false discovery rate set to 0.05.

To visualize the time decoder performance and the posterior probability distribution, we calculated a N x N (N = the number of time bins) matrix to summarize the time decoder prediction outputs. For element (i,j) in the matrix, the value represented the probability of a time bin #i was predicted as time bin #j.

#### Calculation of Absolute Decoding Errors in the Fine-scale Time Analysis

After retrieving the posterior probability distribution of decoding responses (the NxN matrix, where N is the number of bins), we calculated the absolute decoding errors for each time bin, using the following equation: 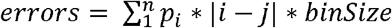, where *n* is the number of bins, *i* are the possible decoder responses, *p*_*i*_ is the posterior probability for response *i*, the ground truth bin index is *j*, and *binSize* is the size of time bin. After obtaining the decoding error curve (as a function of the ground truth bin labels), we fitted the error curve with linear regression. The p values of the slope were reported in the results section.

#### Fine-scale distance decoding analyses

To examine whether continuous distance codes were also present in the scalp EEG power, we discretized data from spatial distance teleportation trials into multiple small “distance” bins and trained SVM classifiers with 2-30Hz power spectra averaged within each distance bin.

To avoid the confounded decoding of fine-scale distance and time, we selected data with only maximal overlap in conceptual distance updating but with zero overlap in the temporal dimension. We selected the 0-2.828s portions of short distance trials, and 2.828s-4.242s portions of long distance trials. While they did not overlap in time ranges, they conceptually covered the same range of spatial distance (see Figure 6A). After the data selection, the 2-30Hz power series of both short and long distance trials were discretized into 11 distance bins, with each distance bin covering 4.42m of distance. For short distance trials, each distance bin occupied 248ms (with a sample rate of 500Hz, 248ms = 124 sampling points), and for long distance trials, each distance bin occupied 248/2 = 124ms (124ms = 62 sampling points). Power estimates within each time bin were averaged. We trained multiclass SVM classifiers with 1280 power spectra features (64 electrodes x 20 frequency). For each classification iteration, 75% of the trials were selected as the training data and 25% of the trials were reserved as the testing data. To increase the independence between training sets and testing sets, a consecutive block of trials was reserved as the testing data, and the rest of data was used for training. We were able to reiterate the classification procedure 37 times. The resulting classification accuracy ratios were averaged across the 37 iterations for each participant, and the 19 participant scores were submitted two-tailed Wilcoxon signed rank tests testing whether they were significantly different than 1.

**Figure 2.**
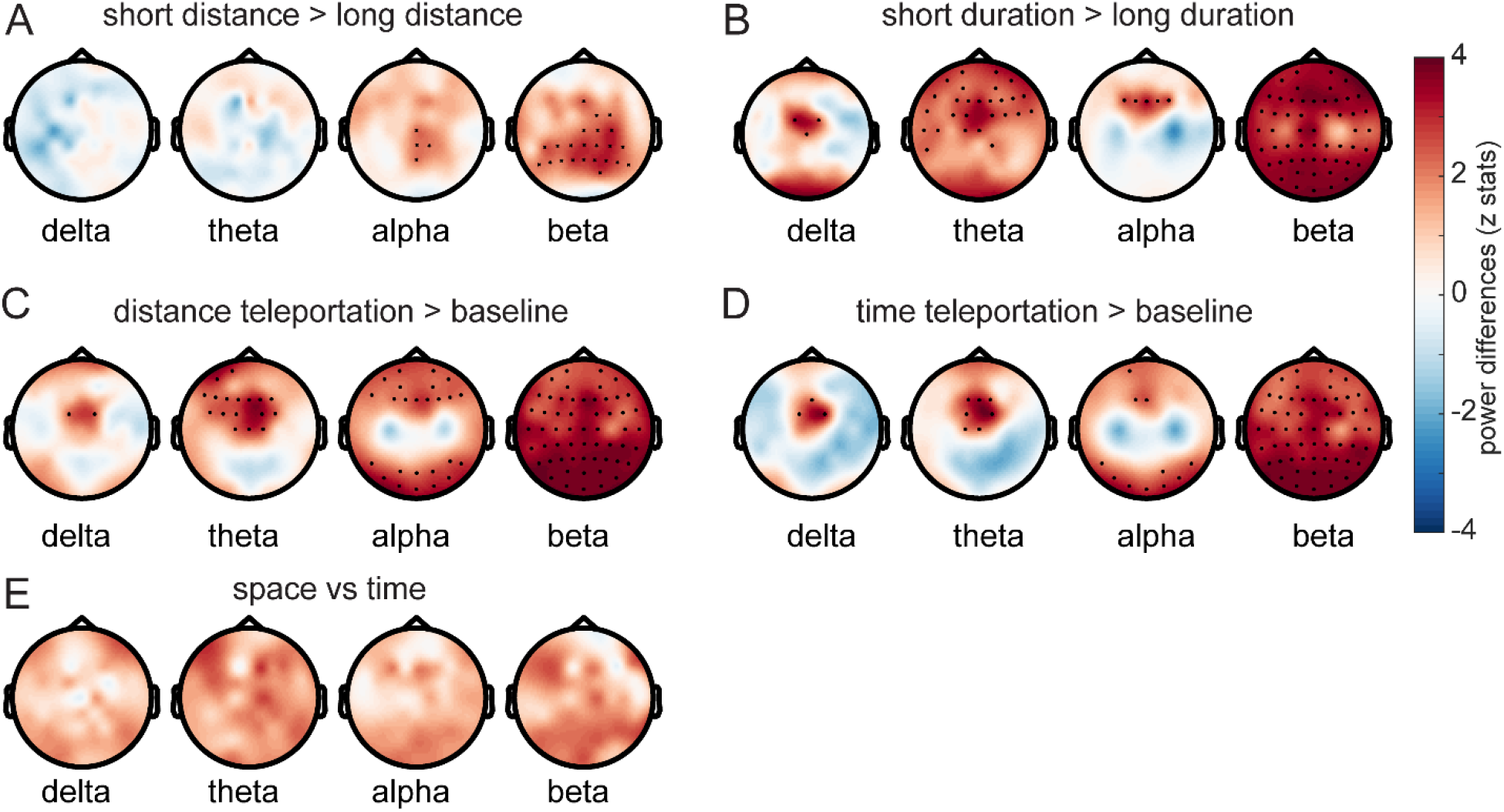
Oscillatory fluctuations present during spatial distance and temporal duration teleportation. **(A)** Short distance teleportation trials resulted in increased alpha and beta power compared to long distance trials. **(B)** Short duration teleportation trials resulted in increased frontal midline delta-theta-alpha power increases, and global beta power increases compared to long duration trials. **(C,D)** Spatiotemporal coding was associated with frontal delta-theta, frontal and posterior alpha, and global beta power increases compared to resting baseline. **(E)** No power differences were observed within the canonical frequency bands between the distance task and the time task. ***Notes***: Black dots are electrodes considered significant after multiple comparison correction. Colors represent the Wilcoxon signed rank tests z statistics.

**Figure 3.**
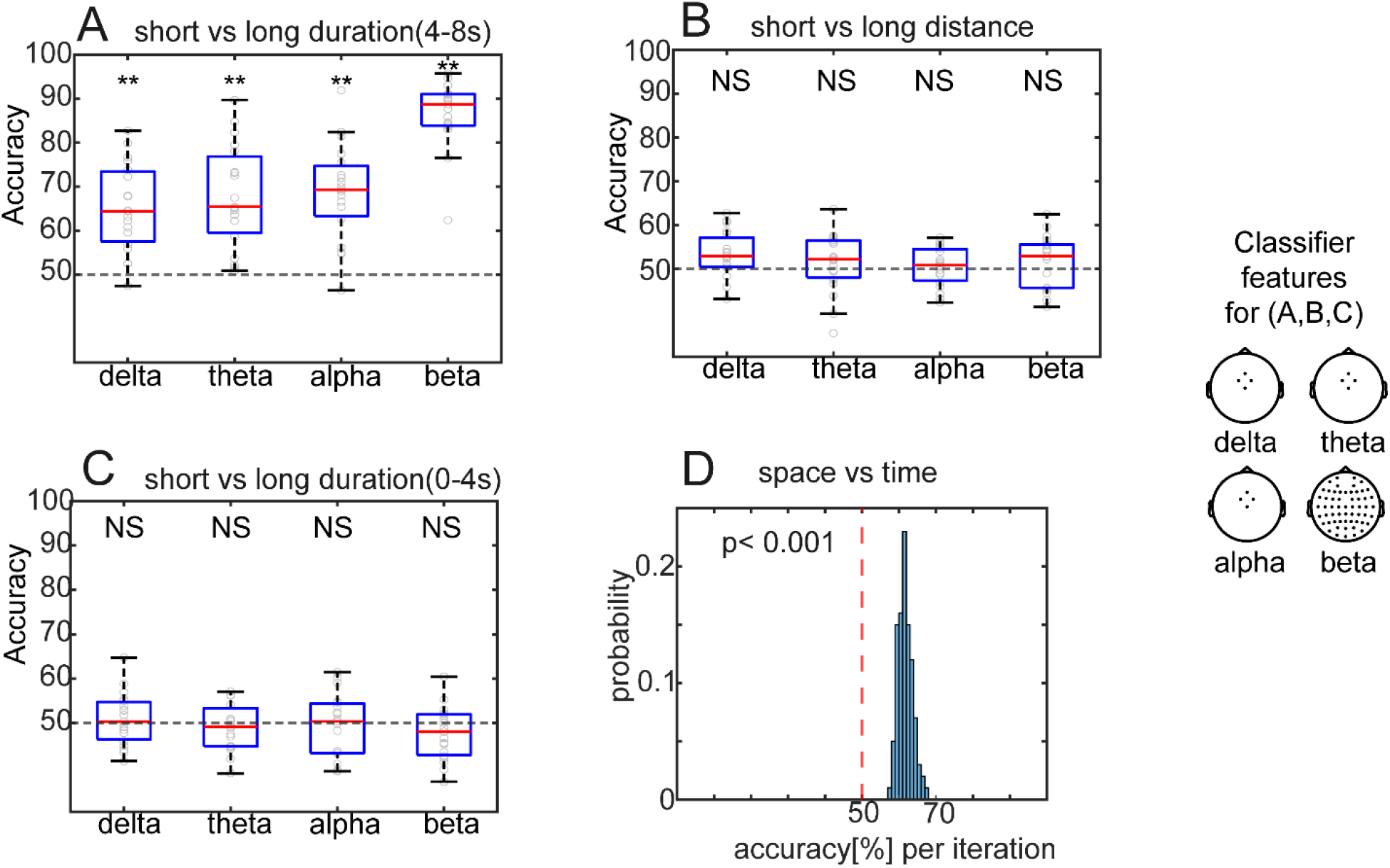
Successful within-task **(A-C)** and between-task **(D)** decoding using power as features. **(A)** Different durations (short vs. long) could be decoded from frontal delta, theta, alpha and global beta power separately. **(B)** Different distances (short vs. long) could not be decoded from frontal midline delta-theta, alpha or global beta power. **(C)** As a control analysis, decoders were not able to differentiate whether a trial was from short duration trials, or from the 0-4s segments of long duration trials. **(D)** When aggregating trials across participants, we were able to decode whether a trial was in the space or time condition based on the single-trial multivariate patterns of power. The histogram of classification accuracies based on 100 iterations is shown**. Notes**: **, all *p*_*FDR*_ = 0.002. Each circle represents a participant in A-C.

**Figure 4.**
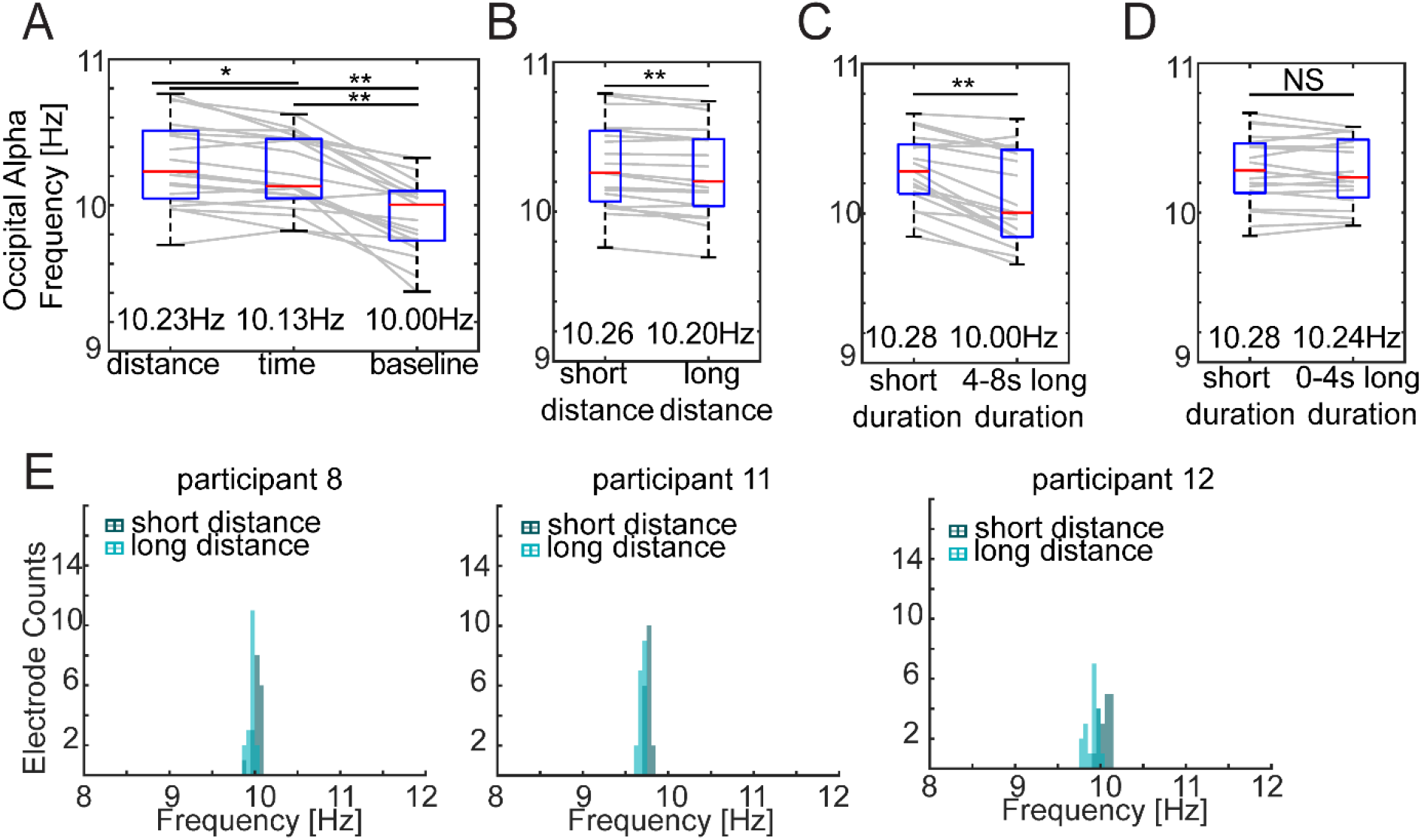
Occipital alpha frequency modulation as a shared mechanism for both spatial and temporal coding. Medians across participants are shown under the box plots. **(A)** The spatial and temporal tasks showed faster alpha frequency than baseline, and the distance task showed faster alpha frequency than the time task. **(B)** In the distance task, traveling a short distance resulted in faster alpha than traveling a long distance. **(C)** In the time task, short duration trials resulted in faster alpha than long duration trials. **(D)** No differences were found between short duration trials and the 0-4s portion of long duration trials. **(E)** Histograms of alpha frequencies at 18 occipital electrodes during the distance task. Data from three example participants were shown. **Notes**: **: *p*_*FDR*_ < 0.01. *: *p*_*FDR*_ < 0.05. NS: not significant.

**Figure 5.**
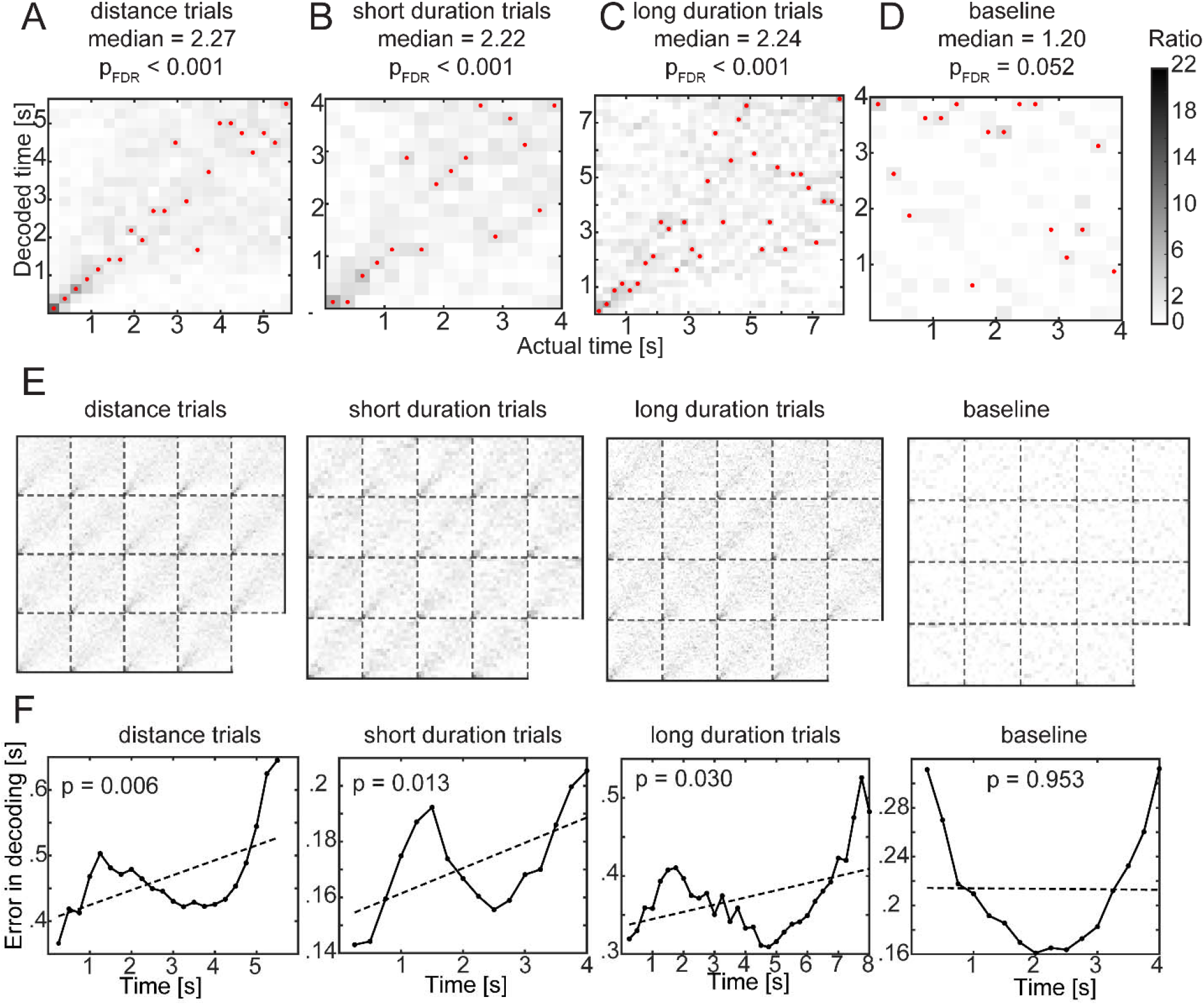
Fine-scale temporal information during the teleportation can be decoded from scalp EEG 2-30 Hz power spectra. Heat maps visualize the posterior probability distribution of the decoder responses. High classification accuracy is indicated by dark colors on the diagonal. **(A,B,C,D)** Fine-scale timing information can be decoded from 2-30 Hz power in the distance task and time task, with accuracies significantly higher than chance level and higher than the baseline task. Medians of accuracy ratios across 19 participants were reported. **(E)** Decoder response probability distributions from 19 participants. Each sub square displays the time decoding heatmap from one participant. **(F)** Decoding errors linearly increased as time progressed in the spatial and temporal tasks, but not in the baseline task. Dashed lines indicate the linear regression fitting models of the decoding errors. **Notes**: Units of the colorbar are accuracy ratios. **Red dots** mark the highest posterior probability in decoder responses.

**Figure 6.**
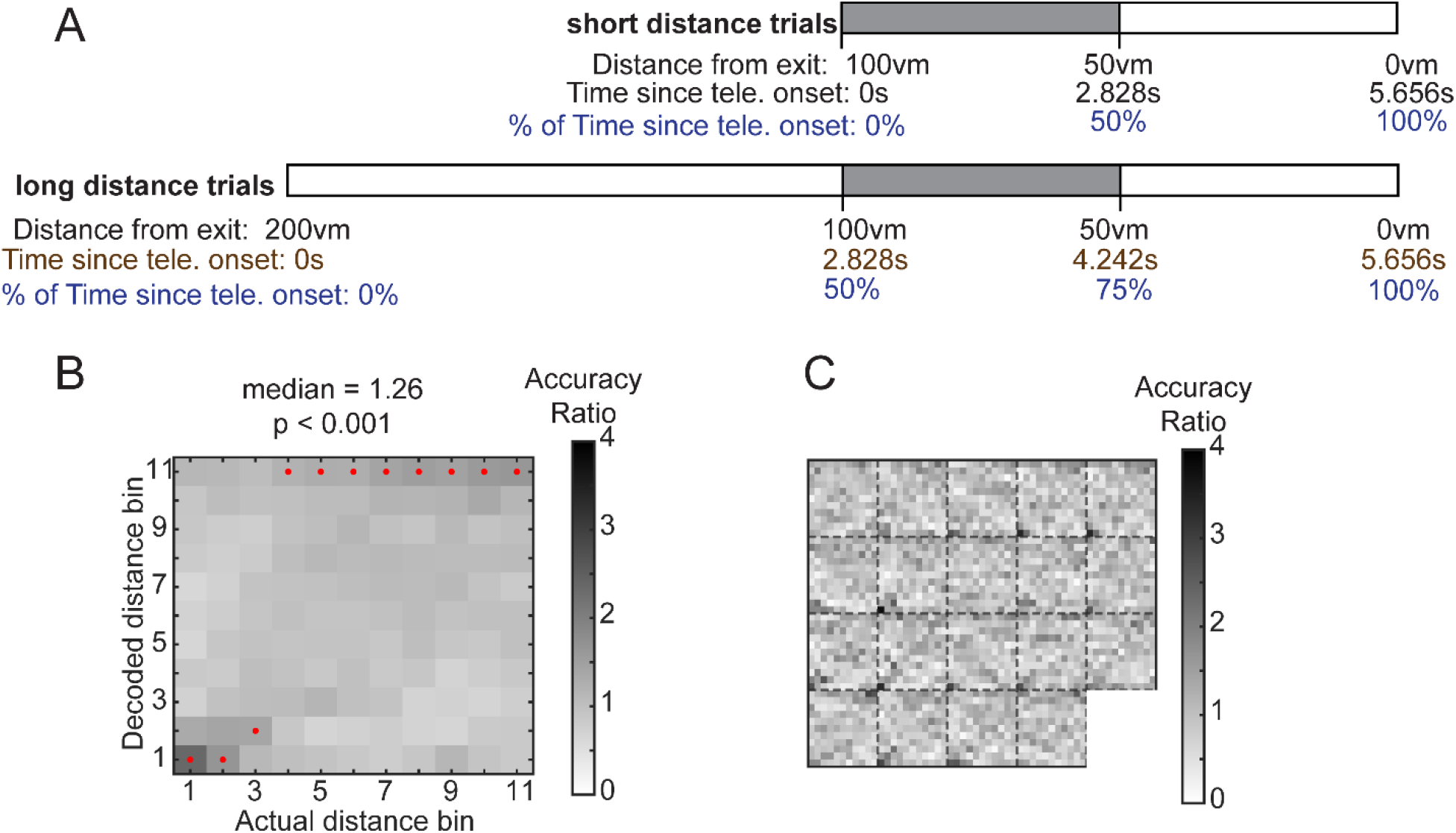
Fine-scale distance information during teleportation can be decoded from multivariate power patterns. Heat maps visualize the posterior probability distribution of the decoder responses. **(A)** Decoding fine-scale distance information while taking care of the temporal confound. To minimize the dependence between temporal and distance information, we selected data (the shaded portions) from both short distance trials and long distance trials that had zero overlaps in the temporal dimension. **(B)** Fine-scale distance information could be decoded in the distance task. **(C)** Posterior probability distributions plotted for each participant. Each sub square displays the distance decoding heatmap from a participant. **Notes**: **Red dots** mark the highest posterior probability in decoder responses.

## Results

### Participants correctly judged spatial and temporal teleportation durations with high accuracy

Participants performed well above chance in both the spatial and temporal teleportation tasks. For the spatial task, out of 48 trials, participants on average made 0.68 errors (SD = 0.89) in judging how far the distance they travelled at the first attempt. For the temporal task, out of 48 trials, participants on average made 1.79 errors (SD = 2.51) in judging how long they spent inside teleporters. On average, participants finished the spatial task within 53.46 (SD = 12.73) minutes and the temporal teleportation task within 52.35 (SD = 9.24) minutes.

### Within-task comparisons: Longer distances traveled associated with decreases in alpha and beta power compared to shorter distance traversals

We first tested the within-task difference hypothesis in the spatial distance task. We compared delta, theta, alpha and beta power between short distance and long distance teleportation trials, and used a cluster-based permutation test for multiple comparison correction. When comparing short distance vs. long distance trials, the permutation test returned a cluster with a p value of 0.015. For short distance trials, we found higher alpha power at central electrodes (Figure 2A, Pz, CP2, Cz, CPz, Cohen’s d: 0.55, averaged log10 alpha power for short distance: [median±SD] = 4.99±0.34, long distance: 4.91±0.32) and higher beta power over central-posterior electrodes (Cohen’s d: 0.91, averaged beta power for short distance: [median±SD] = 4.51±0.26, long distance: 4.50±0.26.) These findings support a possible role for alpha and beta power changes in spatial distance coding.

### Within-task comparisons: Longer temporal durations were associated with frontal delta-theta-alpha power and global beta power decreases compared to shorter temporal durations

We then tested the within-task difference hypothesis for temporal duration teleportation by comparing the power spectra between short duration and long duration trials (Figure 2B). The cluster-based permutation test returned a positive cluster (p < 0.001). This effect was most pronounced over frontal midline electrodes for delta power (Cohen’s d: 1.03, short duration: [median±SD] = 4.47±0.22, long duration: 4.42±0.23), over frontal electrodes for theta power (Cohen’s d: 0.97, short duration: [median±SD] = 4.86±0.19, long duration: 4.83±0.20), and over frontal electrodes for alpha power (Cohen’s d: 0.98, short duration: [median±SD] = 4.35±0.24, long duration: 4.32±0.25). We also found global beta power changes (Cohen’s d: 1.63, short duration: [median±SD] = 4.59±0.25, long duration: 4.55±0.26.)

To further confirm the role of frontal midline theta oscillations in duration timing, we trained a binary classifier to decode types of temporal durations in the teleporter (Figure 3). We successfully decoded whether a trial was a short duration trial or the 4-8 s portion of a long duration trial (Figure 3A, classifiers trained with frontal-midline delta power: [median±SD] = 64.40±9.89%, frontal-midline theta: 65.42±11.68%, frontal-midline alpha: 69.34±10.89%, global beta: 88.76±7.51%, all p_corrected_ = 0.002). However, we could not decode the distance travelled in the teleporter significantly above chance (Figure 3B, classifiers trained with frontal-midline delta power: 52.93±5.09%, p_corrected_ = 0.06, theta: 52.26±6.77%, p_corrected_ = 1, alpha: 50.88±4.47%, p_corrected_ = 1, beta: 52.94±5.95%, p_corrected_ = 1), suggesting frontal midline delta-theta-alpha power, and global beta power alone contained sufficient information regarding the temporal duration being coded but not the distance traveled.

As an additional control analysis, we trained the same classifier with frontal-midline delta-theta-alpha power and global beta power to discriminate the 0-4s portion of the long duration trials from the short duration trials. This served as a control because participants could not have known what types of durations they experienced until they crossed the 4s threshold within the teleporter. Indeed, the classifier was not able to decode whether the trials were short duration trials (4s) or the 0-4s portion of long duration trials (Figure 3C, delta: 50.33±5.88%, theta: 49.16±5.33%, alpha: 50.34±7.25%, beta: 48.04±5.94%, all p_corrected_ > 0.05). Together, these findings support a general role for global beta power changes in spatiotemporal processing, and a unique role of frontal midline delta-theta-alpha oscillations, in coding temporal durations.

### Between task comparisons: Spatial and temporal teleportation did not induce focal differences in delta, theta, alpha, or beta power

To test our between-task hypothesis regarding differences in oscillatory codes between spatial and temporal tasks, we compared the power spectra among spatial, temporal and baseline tasks (Figure 2C, 2D.)

For both contrasts (*distance task* > *baseline; time task* > *baseline*), the cluster-based permutation tests returned a significant positive cluster with p values < 0.001. The effect was most pronounced over frontal midline electrodes for delta power (Cohen’s d for distance vs baseline: 0.60, distance-baseline: [median±SD] = 0.12±0.27, Cohen’s d for time vs baseline: 0.77, time-baseline: [median±SD] = 0.12±0.15), over frontal electrodes for theta power (Cohen’s d for distance vs baseline: 1.04, distance-baseline: [median±SD] = 0.07±0.07, Cohen’s d for time vs baseline: 1.01, time-baseline: [median±SD] = 0.05±0.08), and over frontal and occipital electrodes for alpha power (Cohen’s d for distance vs baseline: 0.82, distance-baseline: [median±SD] = 0.20±0.18, Cohen’s d for time vs baseline: 0.76, time-baseline: [median±SD] = 0.10±0.21). We also found widespread increases in beta power (Cohen’s d for distance vs baseline: 1.81, distance-baseline: [median±SD] = 0.16±0.08, Cohen’s d for time vs baseline: 1.80, time-baseline: [median±SD] = 0.14±0.07). The findings suggest that compared to a passive baseline, participants showed distinct oscillatory profiles while maintaining spatiotemporal information during the teleportation tasks, which was consistent with their high performance in the behavioral tasks.

Next, we asked whether the power spectra profiles differed between the spatial distance and temporal duration task (Figure 2E). The cluster-based permutation test did not reveal any clusters with a p-value lower than threshold. This suggests that the spatial and temporal teleportation tasks did not differ in overall power when compared within each of the canonical frequency bands (delta, theta, alpha and beta bands).

### Between task comparison: Successful decoding of spatial and temporal trials based on single-trial multivariate patterns of power

It could be possible that spatial and temporal coding did not differ in terms of power changes in focal frequency bands; instead, spatiotemporal coding might differ in the multivariate patterns across electrodes and frequencies in a manner that generalizes across participants. To test this possibility, we used multivariate power features to classify whether trials were from the spatial or temporal task. The classifier revealed above chance classification of task labels (Figure 3D, median = 61.46%, SD over 100 iterations = 1.97%, Wilcoxon signed-rank test, z = 8.68, p < 0.001.) These findings suggest the single-trial multivariate patterns significantly differed between spatial and temporal tasks in a manner that generalized across participants. The findings together support a notion of a partially independent space-time code.

### Alpha frequency modulation: A common mechanism for spatial and temporal judgments

We hypothesized that occipital alpha frequency modulation could be an additional form of distance and duration coding in our teleportation task, as suggested by (Cao & Händel, 2019; Samaha & Postle, 2015). To test this idea, we first assayed whether there were differences in occipital alpha frequencies during the teleportation tasks compared to the task-irrelevant resting baseline. Both spatial and temporal teleportation tasks showed faster occipital alpha frequencies than the baseline (Figure 4A, spatial task: [median±SD] = 10.23±0.30Hz, temporal task: 10.13±0.25Hz, baseline: 10.00±0.26Hz; spatial task vs. baseline: Wilcoxon signed rank test, z = 3.74, p_corrected_ = 0.001; temporal task vs. baseline: z = 3.78, p_corrected_ = 0.001.) These findings suggest that occipital alpha frequencies were significantly altered during spatiotemporal coding compared to a resting baseline.

Second, we asked whether occipital alpha frequency differed between the spatial and temporal tasks. Comparing across all participants, the spatial distance task showed significantly faster occipital alpha compared to the temporal teleportation task (Figure 4A, z = 2.62, p_corrected_ = 0.026) The findings of differences in alpha frequencies between spatial and temporal teleportation tasks might reflect another distinction in oscillatory codes for spatiotemporal information.

Therefore, we asked whether the observed occipital alpha frequencies were sensitive to distance and duration information. We first compared the averaged alpha frequency at occipital electrode sites for short vs. long distance trials. When comparing across participants, results revealed that occipital alpha oscillations were of higher frequency for short distance trials compared to long distance trials (Figure 4B, short distance: [median±SD] = 10.26±0.29Hz, long distance: 10.20±0.30Hz, z = 3.38, p_corrected_ = 0.003). Occipital alpha frequency also varied between short and long temporal duration trials. Occipital alpha frequency was faster for short duration trials than the 4-8s portion of long duration trials (Figure 4C, short temporal duration: [median±SD] = 10.28±0.24Hz, long temporal duration(4-8s): 10.00±0.31Hz, z = 3.58, p_corrected_ = 0.002.)

As a control analysis, we tested whether there were differences in occipital alpha frequencies for short duration trials vs. the 0-4s portion of long duration trials. The alpha frequencies did not differ (Figure 4D, short temporal duration: [median±SD] = 10.28±0.24Hz, long temporal duration (0-4s): 10.24±0.23Hz, z = 0.76, p_corrected_ = 1.) Together, these findings support alpha frequency modulation as a shared mechanism for coding spatial distance and temporal durations.

### Fine-scale temporal information was decoded from multivariate patterns of 2-30Hz power spectra

We next tested whether temporal duration codes might be present in the EEG data at a finer scale, inspired by Bright et al. (2020), for example, at the level of 250 milliseconds. Therefore, we trained classifiers on 2-30Hz power to decode times since onset of teleportation. We were able to decode fine-scale temporal information from distance teleportation trials significantly above chance (Figure 5A, accuracy: [median±SD] = 10.34±1.32%, accuracy ratios: [median±SD] = 2.27±0.29%, Wilcoxon signed rank test, z = 3.82, p_corrected_ < 0.001), from short duration trials (Figure 5B, accuracy: [median±SD] = 13.87±1.67%, accuracy ratios: [median±SD] = 2.22±0.27, z = 3.82, p_corrected_ < 0.001), and from the long duration trials as well (Figure5C, accuracy: [median±SD] = 6.99±0.95%, accuracy ratios: [median±SD] = 2.24±0.30, z = 3.82, p_corrected_ < 0.001). As a control analysis, we applied the fine-scale time decoder for data obtained in the baseline task. The decoder was able to decode time from the baseline data marginally better than chance after multiple comparison correction (accuracy: [median±SD] = 7.50±1.87%, accuracy ratios: [median±SD] = 1.20±0.30, z = 2.37, p_corrected_ = 0.052). However, time decoding performance for the baseline task was significantly worse than those in the temporal and distance tasks (baseline < distance task, baseline < short duration trials, baseline < long duration trials: all z = −3.82, p < 0.001.) These findings suggest the intriguing possibility that fine-scaled temporal codes are embedded in low-frequency oscillations.

We note that following entry into the teleporter, participants exhibited a P300-like ERP response (Polich, 2007) at Cz electrode. Therefore, we repeated the fine-scaled time classification analyses, with the grand averaged EEG traces subtracted from every trial. After removing the grand ERP responses, we were still able to successfully decode continuous-like temporal information from the distance teleportation trials (accuracy: [median±SD] = 11.08±1.29%, accuracy ratios: [median±SD] = 2.44±0.28, Wilcoxon signed rank test, z = 3.82, p_corrected_ < 0.001), from the short duration trials (accuracy: [median±SD] = 15.52±1.93%, accuracy ratios: [median±SD] = 2.48±0.31, z = 3.82, p_corrected_ < 0.001), and from the long duration trials (accuracy: [median±SD] = 8.22±1.08%, accuracy ratios: [median±SD] = 2.63±0.35, z = 3.82, p_corrected_ < 0.001.)

Further, to exclude the possible contribution of movement-related in early onsets of a trial, we removed the first second of teleportation epochs and repeated the fine-scale time decoding analyses. We were again able to successfully decode fine-scale time information from distance teleportation trials above chance (accuracy: [median±SD] = 8.80±1.05%, accuracy ratios: [median±SD] = 1.58±0.19, z = 3.82, p_corrected_ < 0.001), from short duration trials (accuracy: [median±SD] = 12.65±1.97%, accuracy ratios: [median±SD] = 1.52±0.24, z = 3.82, p_corrected_ < 0.001), and from long duration trials above chance as well (accuracy: [median±SD] = 5.67±0.89%, accuracy ratios: [median±SD] = 1.59±0.25, z = 3.82, p_corrected_ < 0.001.)

### Decoding errors linearly increased as time progressed forward

We noticed a qualitative pattern that the decoding responses were less precise as time progressed forward in the posterior probability distribution of time decoding responses. To quantitatively test this, we calculated the absolute decoding errors for each timebin and fitted the error curves with a linear regression model (Figure 5F). Results of the linear regression fitting indicated that the decoding errors were significantly larger for later time bins; this effect was found in the distance trials, short duration trials, long duration trials, but not in the baseline task (for distance trials: slope [estimate, standard error (SE)] = [0.02, 0.01], t = 3.03, p = 0.007; for short duration trials: slope [estimate, SE] = [0.01, 0.003], t = 2.70, p = 0.017; for long duration trials: slope [estimate, SE] = [0.01, 0.003], t = 2.56, p = 0.016; for the baseline task: slope [estimate, SE] = [-0.0004, 0.01], t = −0.04, p = 0.97.) The results suggest that the fine-scale temporal information revealed by the decoders are aligned with the human behavioral findings of increased variability for longer reproduced durations (Ivry & Hazeltine, 1995; Rakitin et al., 1998). We discuss the implications in discussion.

### Fine-scale distance information was also present in multivariate patterns of 2-30Hz power

Given our findings with fine-scale temporal information, we also tested whether fine-scale distances could be decoded using the same approach. Indeed, we found that the classifiers were able to decode fine-scale distance information from the spatial task (Figure 6A, accuracy: [median±SD] = 11.45±1.60%, accuracy ratios: [median±SD] = 1.26±0.18, Wilcoxon signed rank test, z = 3.70, p < 0.001). The findings of the fine-scale distance code support the possibility that participants linearly updated their spatial position inside teleporters. The demonstrations of both fine-scale distance and temporal codes in the multivariate power spectra patterns reveal another common aspect that exists in spatiotemporal coding.

## Discussion

In the current study, we tested whether neural oscillations recorded at the scalp supported maintenance of spatial distance and temporal duration information. Decades of research support a role for low-frequency oscillations, both in cortex and hippocampus, in coding spatial information during navigation (McFarland et al., 1975; Vanderwolf, 1969; Kropff et al., 2021; for reviews, see Jacobs, 2013; Watrous et al., 2011). To attempt to disentangle space and time, whose changes are strongly intertwined in movement speed, participants experienced teleportation of different spatial distance and temporal durations in the absence of any optic flow or other sensory input to provide cues about speed, similar to the design in Vass et al. (2016). Results from power spectra analyses suggested the sensitivity of central-posterior alpha power and global beta power for spatial distances, and a role of frontal theta and global beta power changes for temporal duration. Furthermore, the analysis of instantaneous alpha frequencies revealed a robust association between alpha frequency and magnitudes of distances and durations, suggesting alpha frequency modulation as a potential common mechanism for spatial and temporal coding. Classifiers trained on power spectra further support the hypothesis that both distance and temporal information could be decoded from scalp EEG signals at a fine-scale resolution.

Given that hippocampal delta-theta power display a distance code (Bush et al., 2017; Vass et al., 2016), and a connectivity between rodent’s prefrontal and hippocampal theta during mobility (Siapas et al., 2005; Young & McNaughton, 2009), we are surprised to find that the cortical delta-theta power did not exhibit significant differences between short distance and long distance trials. This null finding cannot be explained by the failure of task design, or the absence of spatial coding during the teleportation period. This is because participants demonstrated high accuracy in identifying distances travelled upon exiting the teleporters, and power spectra analyses revealed significantly different oscillatory profiles for the distance task compared to baseline (Figure 2C). What could lead to such a disconnect? Here, we offer three speculations on the null findings linking cortical theta and spatial distance coding. One possibility is that prefrontal theta oscillations are phase locked but not amplitude locked to hippocampal theta (Young & McNaughton, 2009), and therefore phase information in frontal theta but not power changes code spatial distance duration (see Watrous et al., 2013, for an example of this). This is an issue we cannot address in the current study because scalp EEG does not give reliable access to hippocampal signals. A second possibility is that frontal midline theta may be locked to the temporal-processing or memory-related components, but not the movement-related components, of hippocampal (HPC) theta oscillations (Goyal et al., 2020; Watrous et al., 2013). A third possibility is that hippocampal movement-related theta oscillations manifest in the cortex within the traditional alpha band (8-12Hz) consistent with the alpha frequency modulation we observed for both spatial and temporal judgments. The third interpretation is consistent with recent reports (Aghajan et al., 2017; Bohbot et al., 2017; Goyal et al., 2020) that hippocampal movement-related theta oscillations, particularly during real-world movements, manifest most prominently above 8Hz, which would align with the frequency range of traditional alpha band (8-12 Hz) rather than theta band (4-8 Hz).

Our results supporting a role for frontal delta-theta power but not distance coding have important implications. In the power spectra analysis, we found frontal midline delta-theta and frontal alpha power sensitive to the temporal durations, while central-posterior alpha power was sensitive to the distance information. The results provide further evidence for partially independent codes for space and time in the human brain. Our findings demonstrating cortical beta oscillations sensitive to temporal duration align with previous reports of timing-related beta power in time production domain (Grabot et al., 2019; Kononowicz & van Rijn, 2015), and movement-related frontal midline delta-theta increases (Liang et al., 2018). On the other hand, our findings regarding central-occipital alpha oscillations related to distance are consistent with notions that human navigation is enriched with regarding to visual input (Ekstrom, 2015), with occipital alpha oscillations particularly sensitive to visual-related changes (such as optic flow, Cao & Händel, 2019). As proposed by Goyal et al. (2020), a theoretical link might therefore exist between HPC movement-related theta and occipital alpha oscillations. For example, eye closure induces alpha power increases both at occipital sites and in hippocampus (Geller et al., 2014). Our current results would suggest differing roles in navigation for frontal midline theta (4-8 Hz) and occipital alpha (8-12 Hz), which were both found relevant to movement (Liang et al., 2018), and frontal midline theta and occipital alpha oscillations could possibly cooperate to support task-dependent spatial or temporal processing. Therefore, a helpful next step would be to determine how these signals coordinate between hippocampus and cortex in our task using ECoG.

We note that when we compared the power spectra of the spatial and temporal teleportation task, we did not find significant differences. Yet, we were able to classify whether a trial was from the spatial or temporal task with an accuracy better than chance in a manner that was generalizable across participants. This suggests the classifiers captured higher-order differences (perhaps the underlying connectivity patterns) between the oscillatory coding of space and time, other than the mean of power fluctuations. One future direction is to examine the affinity of connectivity patterns for spatial coding and temporal coding, using a similar behavioral task used in this study. We predict that the networks for spatiotemporal coding should diverge, both measured using scalp EEG data, and using intracranial EEG data (as proven by Watrous et al., 2013).

In addition to our findings that spatial distance and temporal duration involve differences in oscillatory codes, both for short vs. long teleportation durations and in their multivariate patterns, we also found a common role for alpha frequency modulation in supporting spatiotemporal coding. Specifically, we found faster occipital alpha for smaller magnitudes of durations/distances. What roles could endogenous alpha frequency modulation possibly play here? One explanation is the processing-speed theory, whereby occipital alpha frequency indexes the processing speed of incoming sensory information (Klimesch et al., 1996). We speculate that the sensory processing speed differed between short and long duration trials because of their different cognitive demands. To complete the temporal task, participants only needed to track time passage in the teleporter up to 4s, and not beyond 4s, and therefore the cognitive demands differed between 0-4s and 4-8s portions of the temporal task.

In contrast to the processing-speed account, another possibility however, relates to a perceptual resolution account. For example, it could be that occipital alpha frequency is linked to the perceptual resolution of duration timing. For example, individuals with 10 Hz resting occipital alpha oscillations might discriminate two temporal durations with a minimum of 100ms (1/10) differences, and those with 12 Hz resting alpha could discriminate two durations with 83.33ms minimal differences (1/12). This perceptual resolution account is also supported by Samaha and Postle (2015) showing that occipital alpha frequency reflects the “refresh rate” of visual perception and occipital alpha represents the perceptual unit of temporal processing (Cecere et al., 2015). Future studies should investigate the potential causal links between occipital alpha frequency and spatiotemporal processing, given recent findings that tACS-induced alpha frequency shifts led to shifts in subjective time experiences (Mioni et al., 2020) and that clinical Alzheimer populations show irregularities in parietal alpha oscillations (Montez et al., 2009).

Given that we found alpha frequency modulation and beta power fluctuations related to both spatial and temporal judgments, our results also provide evidence for a common mechanism for spatial and temporal coding involving magnitude estimation. Although distance-related beta power has rarely been studied in a scalp EEG setting, the timing-related beta power we observed has been noted in predicting the accuracy and precision of time production (Grabot et al., 2019; Kononowicz & van Rijn, 2015). Our findings suggest that beta oscillations may reflect a common magnitude representation underlying both spatial and temporal processing, and that such distance and fine-scale temporal information could be widely accessible in neocortical regions, including early sensory and motor cortices. Future studies can bridge the gap of research between spatial and temporal processing, and further elaborate the roles of beta oscillations in spatial coding vs. temporal coding, with a variety of tasks such as estimating and reproducing spatial distance with a path integration task (Harootonian et al., 2020).

Another important finding from our study is the ability to decode fine-scale distance and temporal information from cortical low-frequency power spectra. Interestingly, when attempting to decode temporal information, we showed that the decoding error linearly increased as the time bins progressed forward. These findings are closely aligned with the behavioral findings in which humans show larger variability in time reproduction responses for longer intervals (Ivry & Hazeltine, 1995; Rakitin et al., 1998). One intriguing possibility is that the cortical low-frequency oscillations support a fine-scale representation of temporal intervals. Future studies can test this possibility by linking the decodability of fine-scale time information and the accuracy/precision of time reproduction in human participants.

Notably, our findings of decodable fine-scale temporal information are qualitatively similar to the findings done with entorhinal temporal context cells (Bright et al., 2020). The tenet of a unified math model of space and time (Howard et al., 2014) is that the neural representations are the Laplace transform of space and time, coded through the exponentially decayed firing rates of neurons. However, the theory does not directly predict or rule out the involvement of neural oscillations in coding space and time. Here we demonstrated that neural oscillations could yield a similar time representation possibly with scale invariance, and we suggest that neural oscillations could be a synergistic component on top of single neuron firing rates for spatiotemporal coding. Another question that should be clarified through future studies is whether the neural representations of spatial distance also possess scale invariance like the representations of time (i.e., reproducing longer distances are associated with greater variability in responses.) Behavioral findings suggest path integration errors systematically scaled with path lengths (Harootonian et al., 2020), which will predict linearly increases in decoding errors as distances increase. Future studies should further test the links between oscillatory representations of fine-scale space and time, and the behavioral phenomena of spatiotemporal reproduction, using a reproduction paradigm, such as reproducing space and time in virtual reality (E. M. Robinson & Wiener, 2020).

### Limitations

It is worth considering some potential limitations with our paradigm which we nonetheless believe do not undermine or challenge our findings. One concern could be that because participants knew how far they would travel before entering the teleporter, distance coding was therefore transient and completed *before* entering the teleporters, thus nullifying the existence of distance coding *during* the teleportation. We note, however, that maintenance of distance information *during* the teleportation was still necessary for accurate performance in the spatial teleportation task. When participants entered the teleporter, while they knew beforehand whether it was a short or long distance, they had to maintain this information during teleportation to make the correct decision upon exiting the teleporter. Our interpretation of perceiving spatial distance prior to decisions about movement is consistent with a rich literature in human spatial navigation, suggesting that humans first estimate distance based on perceptual cues and then attempt to maintain this in working memory as they actively navigate to different goals (Knapp & Loomis, 2004; Philbeck et al., 1997; Philbeck & Loomis, 1997). Using a similar spatial distance teleportation design, Vass et al. (2016) showed that the spatial distance teleportation task resulted in different oscillatory profiles from those during the resting state (viewing a laptop black screen outside the experimental context). We similarly found a clear difference between teleportation and a resting baseline task. These findings suggest that the spatial teleportation task triggered distance information processing absent in a resting state condition.

Another concern could be that movement-related noise from the navigation phase permeated into the EEG data during the teleportation, thus confounding the findings we presented here. Note that the amount of noise, if any, should be identical between short and long trials, and between the spatial and temporal tasks, given that participants stood still after they entered the teleporter. Therefore, noise should not confound the findings regarding the contrasts of EEG responses between short and long trials, or between the spatial and temporal tasks.

## Conclusions

Our study addressed an important issue regarding whether spatial and temporal processing share common or distinct mechanisms (Eichenbaum & Cohen, 2014; Ekstrom et al., 2011; Frassinetti et al., 2009; Gauthier et al., 2019, 2020; Watrous et al., 2013). Our findings suggest that spatial and temporal judgments during navigation differ as a function of power changes within specific frequency bands: while spatial judgments resulted in changes in cortical alpha and beta power, while different temporal durations were linked to changes in frontal midline delta-theta, frontal and posterior alpha, and global beta power. Consistent with the idea of separable representations for space and time, spatial and temporal discounting are behaviorally distinctive from each other (E. Robinson et al., 2019), estimating spatial distance are subject to large errors (Zhao, 2018) while estimating suprasecond durations can be performed with high accuracy (Grabot et al., 2019), and spatial and temporal estimation errors distort in opposing manners (Brunec et al., 2017). Previous reports have also hinted at a dissociation between space and time at the neural level although using different paradigms in which temporal information, in particular, involved order and not duration (Ekstrom et al., 2011; Watrous et al., 2013). More generally, evidence exists for and against the notion that space and time processing are of the same nature, and we also found evidence for alpha frequency modulation as a common mechanism for spatial and temporal coding. Thus, one implication of our study is that there are both distinct and common mechanisms related to how we process spatial distance and temporal durations.

## Acknowledgements

This research was supported by National Science Foundation (NSF BCS-1630296, A. D. E.). We thank Stephanie Doner for the assistance in scalp EEG data collection, Eva Robinson for feedbacks on the manuscript, and the participants for being part of this study.

